# Mitoxantrone Hydrochloride Targets APP and LRRK2 to Improve Neurodegeneration in Parkinson’s Models

**DOI:** 10.1101/2025.10.06.680832

**Authors:** Haitao Tu, Zhi-Wei Zhang, Chaitanya K. Jaladanki, Muhammad Yaaseen Gulam Mohamed, Wuan-Ting Saw, Sook-Yoong Chia, Yinxia Chao, Zhidong Zhou, Zhong Pei, Hao Fan, Eng-King Tan, Li Zeng

## Abstract

**Background:** Parkinson’s disease (PD) is the second common neurodegenerative disorder, driven by the loss of dopaminergic neurons and pathological α-synuclein protein accumulation. Currently, there is no disease-modifying therapy that can halt PD progression. Our previous study uncovered a critical pathogenic feed-forward loop between amyloid precursor protein (APP) and leucine-rich repeat kinase 2 (LRRK2), in which the two proteins mutually enhance each other’s expression, ultimately leading to mitochondrial dysfunction and neurotoxicity. Targeting this vicious cycle represents a promising therapeutic strategy for PD.

**Methods:** To discover novel inhibitors targeting this axis, we performed high-throughput screening of an FDA-approved drug library using a fluorescence-based biosensor system. We identified Mitoxantrone hydrochloride (MH), an antineoplastic agent, as a lead compound that inhibits both APP and LRRK2 expression. Its efficacy was validated in cellular models, including patient induced pluripotent stem cell (iPSC)-derived dopaminergic neurons and human peripheral blood mononuclear cells (PBMCs). Motor behavioural and safety assessments were subsequently conducted in PD mouse models.

**Results:** We demonstrated that MH suppresses both APP and LRRK2 expression in various cell types in a dosage-dependent manner. In addition, MH also inhibits phosphorylation of LRRK2 and its downstream substrate Rab10. We further showed MH inhibits LRRK2 activity through direct binding to its kinase domain. Critically, MH treatment rescued dopaminergic neuron loss and reversed motor deficits in both 6-Hydroxydopamine (6-OHDA)-induced and LRRK2^G2019S^ genetic PD mouse models. Moreover, we found that oral administration of MH is therapeutically effective, providing superior neuroprotection and behavioral recovery without detectable cardiotoxicity or gastrointestinal damage.

**Conclusions:** Our findings demonstrate MH as a compelling, repurposable therapeutic candidate capable of disrupting a core pathogenic mechanism in PD.

## BACKGROUND

Parkinson’s disease (PD) is the second most common neurodegenerative disorder after Alzheimer’s disease (AD). Patients with PD may experience uncontrolled tremor, muscular rigidity, bradykinesia, and postural instability, with cognitive decline in most cases. Pathologically, PD is characterized by the presence of misfolded alpha-synuclein (αSyn) in Lewy bodies, causing a progressive loss of dopaminergic (DA) neurons in the substantia nigra (SN) [1–3]. PD affects approximately 1–2% of individuals over the age of 65, corresponding to nearly 10 million people worldwide [4]. In Singapore, the prevalence is about 3 per 1,000 individuals over the age of 50, and the incidence increases with age [5]. The economic impact is substantial, with the annual healthcare cost for PD patients in the United States estimated at US $52 billion in 2020, and this figure continues to rise, imposing a considerable burden on caregivers and society [6, 7].

To date, there is no disease-modifying therapy for PD, which may partly reflect the complex and incompletely understood pathogenesis involving genetic factors, environmental toxins, and oxidative stress [8]. Current treatments primarily provide symptomatic relief but do not slow disease progression [9]. The loss of dopaminergic neurons in the substantia nigra pars compacta (SNpc) is widely believed to underlie the motor and non-motor manifestations of PD. Dopamine replacement therapies can alleviate symptoms but do not halt neurodegeneration [9, 10]. Over the past two decades, several genes have been implicated in PD, including *SNCA*, *PRKN* (Parkin), *PARK7* (DJ-1), *PINK1*, and *LRRK2*, highlighting the genetic contributions to disease susceptibility and pathophysiology. Among genetic risk factors, pathogenic mutations in *LRRK2* represent the most common contributor, accounting for 5-15% of familial cases in many populations [5].

Inhibition of LRRK2 kinase activity represents a promising therapeutic strategy for PD. Preclinical studies in animal and cellular models indicate that LRRK2 inhibitors can alleviate neurodegenerative pathology and improve motor outcomes [11, 12]. Early-phase clinical trials, including those evaluating DNL201 and DNL151 (BIIB122), have reported favorable safety profiles and adequate brain penetration [13, 14]. Ongoing Phase 2 trials are designed to determine the efficacy of these agents in reducing PD symptoms and slowing disease progression [15, 16]. Nevertheless, important challenges remain, including the need to achieve highly selective inhibition and to establish long-term safety [17]. Although LRRK2 inhibition offers potential benefits such as neuroprotection and attenuation of α-synuclein pathology [18], concerns regarding off-target effects and the incomplete understanding of LRRK2’s physiological functions warrant careful evaluation [19]. Despite these uncertainties, LRRK2 inhibitors continue to represent one of the most promising avenues for the development of disease-modifying therapies in PD.

Our previous studies revealed that mutant LRRK2^G2019S^ enhances processing of amyloid precursor protein (APP) into its transcriptionally active form, the APP intracellular domain (AICD), and that *LRRK2* and APP/AICD form a self-perpetuating pathogenic cycle [20, 21]. This cycle drives excessive mitophagy, mitochondrial dysfunction, α-synuclein accumulation, and decreased tyrosine hydroxylase, culminating in dopaminergic neuron loss and neurodegeneration [20, 21]. Notably, pharmaceutically inhibition of AICD by itanapraced blocks AICD nuclear translocation, disrupting this feed-forward loop and restoring dopaminergic neurons [21, 22]. These findings establish a reciprocal link between *LRRK2* and APP/AICD, highlighting both as promising therapeutic targets for PD and motivating the development of novel, patentable inhibitors.

In this study, to identify novel and highly efficacious inhibitors of APP and *LRRK2*, we employed our in-house fluorescence-based biosensor system to perform high-throughput screening (HTS) of small molecules from FDA-approved libraries. Based on in vitro cell-based assays, Mitoxantrone hydrochloride (MH) emerged as the most potent compound in suppressing both APP and *LRRK2* expression and activity. In silico predictions, followed by cellular validation, indicated that MH inhibits *LRRK2* activity through direct binding to its kinase domain, in addition to its effects via the APP-*LRRK2* feed-forward loop. Notably, MH mitigated dopaminergic neuron loss and reversed motor deficits in a PD mouse model with minimal side effects. These findings highlight the potential of repurposing an FDA-approved drug to target PD through a novel pathophysiological pathway.

## MATERIALS AND METHODS

### Study design

Our previous research established a pathophysiological connection between AD and PD, the two most common neurodegenerative disorders, through the APP-LRRK2 loop [20, 21]. Pharmacological inhibition of *LRRK2* via this loop using the AICD inhibitor itanapraced mitigated neurotoxicity and motor deficits in vitro and in vivo [21]. However, itanapraced required relatively high effective concentrations, 40 µM in vitro and 30 mg/kg in vivo, which are often considered less physiologically relevant [23, 24]. Consequently, we aimed to develop a more potent and physiologically relevant drug targeting the APP-LRRK2 loop for the treatment of PD. Drug repurposing offers a fast, cost-effective approach to identify new treatments for neurodegenerative diseases. Using high-throughput screening of FDA-approved drug libraries allows rapid discovery of candidates with established safety and pharmacokinetic profiles. Many repurposed drugs have pleiotropic effects, such as anti-inflammatory, antioxidant, or mitochondrial protection, making them especially relevant to the complex pathology of disorders like AD and PD [25].

By conducting high-throughput screening for APP transcriptional inhibitors from a library of 1,971 FDA-approved drugs, we identified MH as a lead compound. To evaluate its inhibitory efficacy, we tested MH across multiple cellular models, including HEK293 cells, SH-SY5Y neuroblastoma cells, LRRK2^G2019S^ patient iPSC-derived dopaminergic neurons, and human PBMCs. To further assess the functional impact of MH in PD pathology, we administered the drug to 6-OHDA toxin-induced PD mice and LRRK2^G2019S^ transgenic mice via intraperitoneal injection and oral feeding. Behavioral, biochemical, and toxicity analyses were performed to evaluate therapeutic effects. This study was reviewed and approved by the Institutional Review Board and the Institutional Animal Care and Use Committee (A25006).

### Cell culture

Human neuroblastoma cell line SH-SY5Y (ATCC, #CRL-2266) and HEK293T cell line (ATCC, #CRL-11268) were maintained in Dulbecco’s Modified Eagle Medium (Gibco) supplemented with 10% Fetal Bovine Serum (Gibco) 1% L-GlutaMAX (Gibco), and 1% penicillin-streptomycin and incubated at 37°C with 5% CO_2_. Plasmid transfection in SH-SY5Y and HEK293T was conducted using Lipofectamine® LTX transfection reagent (Invitrogen) following the manufacturer’s instruction. Transfected cells were used for immunoblotting or immunocytochemical staining assays 2 days post-transfection.

### Mouse lines

C57BL/6J mice were obtained from the Animal Holding Units of Lee Kong Chian School of Medicine, Nanyang Technological University, Singapore. The C57BL/6J BAC *LRRK2^G2019S^* transgenic mice were obtained from Jackson Laboratories (#012467). Animals were maintained in accordance with institutional guidelines, and all protocols were approved by the Institutional Animal Care and Use Committee of the National Neuroscience Institute and Tan Tock Seng Hospital. Mice were maintained in a pathogen-free facility and exposed to a 12-hour light/dark cycle. Food and water were continuously available. Male mice were used for all studies.

### High throughput screening (HTS)

pCopGFP-BSA-pAPP–transfected SH-SY5Y cells (15,000 cells per well) were seeded into 96-well plates (Thermo Scientific, #165305) and treated with 10 µM of each compound from the FDA-approved drug library (ApexBio, #L1021). CopGFP fluorescence signals were measured at 0-, 2-, 24-, and 48-hours post-treatment using a plate reader (Tecan, M200). Compounds that reduced CopGFP fluorescence were shortlisted as potential APP transcriptional inhibitors. These candidates were subsequently subjected to a second-round screening using serial concentrations (0.1, 0.5, 2, and 10 µM) following the same procedure. For hit validation, pCopGFP-BSA-pAPP–transfected SH-SY5Y cells (50,000 cells per coverslip) were seeded on poly-L-lysine–coated coverslips (Sigma, #P8920) and treated with varying concentrations of MH for 24 hours. The cells were fixed in 4% paraformaldehyde for 15 minutes at room temperature, counterstained with DAPI (4′,6-diamidino-2-phenylindole) for 15 minutes, and imaged using a confocal microscope (Olympus FV3000).

### Drug MH treatment

MH was purchased from Selleck Chemicals (#S2485) and dissolved in DMSO. Cells were treated with MH for 24 hours, while PBMCs were treated for 1 hour. 6-OHDA PD mouse model was generated as described [21], and MH was administered via i.p. injection (0.5 or 1mg/kg/day for 5 days, total dose=2.5 or 5 mg/kg) or oral gavage (1 mg/kg/day for 5 days, total dose=5 mg/kg) from day 8 to day 13 before behavioural tests.

### Isolation of human PBMCs

The use of human blood from healthy volunteers was reviewed and approved by the Institutional Review Board. Whole blood (10 mL per individual) was collected into EDTA-containing tubes and centrifuged at 500 × *g* for 10 minutes. The plasma layer was carefully removed, and the remaining blood fraction was diluted 1:1 with phosphate-buffered saline (PBS, 1×). The diluted blood was gently layered onto Ficoll-Paque™ PLUS (Cytiva, # 17144002) in a 15 mL conical tube and centrifuged at 400 × *g* for 30 minutes at room temperature with acceleration and deceleration settings of 4. The PBMC layer at the plasma–Ficoll interface was collected and washed once in PBS (400 × *g*, 15 minutes). Red blood cells were lysed using Red Blood Cell Lysis Buffer (Thermo Fisher Scientific, #00-4300-54), followed by two additional PBS washes. The final PBMC pellet was resuspended in appropriate culture medium for subsequent experiments.

### Binding Site Prediction and Receptor Grid Generation

The crystal structures of human LRRK2 (PDB ID:8FO7) were obtained from the Protein Data Bank (PDB). These protein structures were prepared using Protein PrepWizard in the Schrödinger Suite, including assignment of bond orders, addition of hydrogens, optimization of hydrogen-bonding networks, and restrained minimization using the OPLS3e force field [26, 27]. The compound MH was prepared using Ligprep to generate possible ionization and tautomeric states at physiological pH. The geometry was minimized using the OPLS3e force field prior to docking. Potential ligand-binding pockets on LRRK2 were predicted using SiteMap, which assesses pocket properties including volume, exposure, hydrophobic/hydrophilic balance, and hydrogen-bonding capacity. Receptor grids were generated at the centroid of the top-ranked predicted sites using the Receptor Grid Generation module in Glide [28]. The inner and outer grid boxes were set to 15 Å and 20 Å, respectively, to define the docking search region.

### Molecular Docking

Molecular docking was performed using the Glide module of the Schrödinger Suite in standard precision (SP) mode. The OPLS3e force field was used throughout the calculations [27]. Docking scores were employed to rank ligand binding affinities across the identified binding pockets.

### Residue Scanning and MM-GBSA Binding Energy Calculations

Residue-level contributions to MH binding were evaluated using the Residue Scanning module in Schrödinger [27]. All residues within 5.0 Å of MH in the docked complex were individually mutated to other 19 amino acids to assess their influence on binding. The resulting mutant structures were optimised, and changes in binding free energy (ΔΔG) were estimated using MM-GBSA calculations [29]. Residues exhibiting significant destabilization upon mutation were interpreted as essential contributors to ligand stabilization and overall complex integrity.

### Mouse behavioural studies

#### Rotation test

A video recorder is set up on a tripod facing downward above the laboratory floor. Each mouse is placed in standard laboratory beakers and allowed to explore environment for 20 min. Mice are injected with 5 mg/kg amphetamines (i.p.) and placed back into the beaker. The investigator will then record peak rotation over 20 min immediately following apomorphine injection and 20 min after amphetamine injection. These recordings then can be analysed post hoc and full rotations can be counted using fast-forward video playback reporting net rotations over the period (No. of clockwise rotations –No. of anticlockwise rotations).

#### Rotarod test

The Rotarod test measures coordination and motor skill learning, as described [30]. Mice were placed on a barrel-shaped platform that rotated slowly at first (4 rpm), which then gradually accelerated over 5 min (to 40 rpm). Each mouse had to balance for as long as possible before falling off. There is a pressure plate below the apparatus, which stops the time recording when the mouse has activated it. The time taken for each mouse to fall was measured here. The longer the duration, the better the coordination and motor skill learning.

#### Cylinder test

This cylinder test measures the vertical movement of mice. Each mouse was placed in an empty cylinder and a 5-minute timer was simultaneously started. When mice are placed in a novel environment, they tend to stand on their hind legs to better smell the surroundings (a behaviour known as rearing). In this test, the number of rears were recorded for each mouse.

#### Pole test

The pole test is a behavioural test that measures coordination and motor function in mice. A 50 cm pole with a stand was placed in the cage. Extra bedding was placed in the cage to protect the falling mice from the pole. The pole test was performed on 3 consecutive days. For the first 2 days, mice were placed on top of the pole and trained to walk down the pole. The training can be performed a few times until mice can walk down the pole by themselves. For the third day, the timings of each mouse walking down the pole for two rounds were recorded for analysis.

### Plasmids construction

#### pCopGFP-BSD-pAPP

A puromycin (PURO) expression cassette, a blasticidin expression cassette (BSD), a CopGFP expression cassette, and the mCherry driven by a 1463 bp human APP promoter were cloned into the pGL3-basic vector (Addgene, #212936). Vector sequence was confirmed by Sanger sequencing.

#### LRRK2-KD-MT

Mutagenesis overlapping primer sets for LRRK2 A1904E (GCT to GAA) and T2035A (ACA to GCA) were synthesized (IDT) and amplified using 2XMyc-LRRK2-Kinase (Addgene, plasmid # 25071) as the template for overlapping poly chain reaction (PCR).

### Sirius Red staining

Sirius red staining of heart tissue was performed using the Picro-Sirius Red Stain Kit (Cardiac Muscle) (Abcam, #ab245887) following manufacturer’s instruction. Briefly, cryosectioned tissue slides were rinsed in PBS and stained with hematoxylin to visualize nuclei. The sections were then stained with the Sirius Red solution (Abcam, #ab245887) for 60 minutes at room temperature. After staining, slides were briefly rinsed in 0.5% acetic acid for 1 minute, repeated twice, to remove unbound dye and minimize background. The sections were subsequently dehydrated through a graded ethanol series (75%, 95%, and 100%), air-dried, and mounted with coverslips using a mounting medium. Images were acquired using an Olympus microscope.

### Western blot analysis

Western blot analysis was performed as previously described [31]. Briefly, cells were lysed in RIPA buffer (50 mM Tris-HCl pH 7.4, 150 mM NaCl, 2 mM Na2 EDTA, 1% NP-40, 0.1% SDS, 0.5% sodium deoxycholate) supplemented with protease inhibitor cocktail (MedChemExpress, #HY-K0010) and phosphatase inhibitor cocktail (MedChemExpress, #HY-K0022). 20 µg of protein lysates were separated on SDS-PAGE gels and transferred to polyvinylidene difluoride (PVDF) membranes. The blots were blocked with TBST (tris-buffered saline–Tween 20) supplemented with 5% skim milk and incubated with primary antibodies overnight at 4°C. The primary antibodies used in this study were LRRK2 (Abcam, #ab133474), APP, LC3II (Abcam, #ab243506), tyrosine hydroxylase (TH) (Millipore, #MAB318), p62 (Cell Signaling, #39749,), p-α-synuclein (S129) (Cell Signaling, #23706S), cleaved caspase 3 (Cell Signaling, #9661S), pS199/202-Tau (Milipore, #ab9674), α-synuclein (BD Transduction Lab, #610787), pThr^73^ Rab10 (Abcam, #ab230261), Rab 10 (Cell Signaling, #8127,) and β-actin (Santa Cruz, #AC-15). The membranes were washed in TBST, and then incubated with secondary horseradish peroxidase (HRP)-conjugated antibody against mouse IgG HRP (GE Healthcare, #NA931V) and rabbit IgG HRP (GE Healthcare, #NA934V) and developed using an ECL detection kit and captured with a ChemiDoc Imaging System. The images were then quantified with ImageJ and plotted with GraphPad Prism for graphs.

### Immunohistochemistry (IHC) staining

Mice were intraperitoneally anesthetized with ketamine/xylazine (100/10 mg/kg; 0.1 mL/g body weight) and perfused transcardially with 0.1 M PBS (pH 7.4). Brains were collected, fixed in 4% paraformaldehyde at 4 °C overnight, and cryoprotected in 30% sucrose until they sank. Fixed tissues were sectioned at 25 µm thickness using a cryostat (Leica). Sections were washed in PBS and blocked for 30 minutes at room temperature in blocking buffer (1% bovine serum albumin (BSA) and 0.1% Triton X-100 in PBS). Samples were then incubated overnight at 4 °C with primary antibodies against tyrosine hydroxylase (TH; Millipore, #AB9702) and Ki67 (Abcam, #ab15580). After three PBS washes, sections were incubated for 2 hours at room temperature in the dark with appropriate Alexa Fluor–conjugated secondary antibodies: Goat anti-Rabbit IgG Alexa Fluor 488 (Thermo Fisher, #A11034), Donkey anti-Mouse IgG Alexa Fluor 555 (Thermo Fisher, #A31570), and Goat anti-Rabbit IgG Alexa Fluor 647 (Thermo Fisher, #A21245). Nuclear counterstaining was performed with DAPI (Sigma-Aldrich, #B2261). Stained sections were mounted with anti-fade mounting medium onto microscope slides and imaged using a confocal laser scanning microscope (Olympus FV3000).

### Statistical analyses

For each experiment, at least three independent measurements were performed. All statistical analyses were performed using GraphPad Prism 8 software. Data were presented as mean ± SD for all statistical analyses. For 2 group comparison, a two-way Student’s *t* test was used to compare the difference between the 2 groups. For multiple group comparisons under one experimental condition, one-way ANOVA with Tukey’s post *hoc* test was used to compare the differences between groups. For multiple group comparisons under two experimental conditions, two-way ANOVA with multiple comparison was used. The statistical significance levels were set at \**P* < 0.05, \*\**P* < 0.01, \*\*\**P* < 0.001, and \*\*\*\**P* < 0.0001.

## RESULTS

### High-throughput screening identifies MH as an inhibitor of both APP and LRRK2

Our previous studies identified the APP/AICD–LRRK2 loop as a novel pathogenic mechanism in PD, and demonstrated that inhibition of AICD confers neuroprotection and rescues motor deficits in PD models [20, 21]. Based on these findings, we hypothesized that disrupting the APP–LRRK2 vicious cycle may exert therapeutic benefits. Accordingly, we aimed to develop more potent inhibitors targeting the APP–LRRK2 loop for the treatment of PD. Drug repurposing has become an attractive strategy for neurodegenerative diseases, supported by both recent trends and clear therapeutic benefits. Screening FDA-approved drug libraries through traditional high-throughput assays offers a rapid and cost-effective means to identify candidates with established safety and pharmacokinetic profiles [32, 33]. This approach reduces the time, expense, and risk of failure associated with de novo drug discovery, while facilitating faster clinical translation. Moreover, many repurposed drugs exert pleiotropic actions, such as anti-inflammatory, antioxidant, or mitochondrial protective effects, that are particularly relevant to the complex pathology of [25].

To facilitate high-throughput drug screening, we first generated a fluorescence-based reporter vector in which the human APP promoter drives the expression of CopGFP. The fluorescence intensity of CopGFP, monitored by fluorescence microscopy or a plate reader, reflects the expression level of APP in cells. Using this fluorescence-based approach, we screened a drug library containing 1,971 FDA-approved compounds in neuroblastoma SH-SY5Y cells (**Fig. 1A**). After several rounds of screening, shortlisted candidates were further validated for their effects on APP expression by fluorescent imaging and immunoblotting (**Fig. 1A**). From this screening, we identified Mitoxantrone hydrochloride (MH), an anthracycline drug clinically used for the treatment of advanced prostate cancer, acute non-lymphocytic leukemia, and multiple sclerosis, as the lead compound that inhibits APP expression. MH suppressed CopGFP fluorescence intensity in a dose-dependent manner, as measured by a plate reader. At 10 µM, MH significantly reduced CopGFP intensity to 12.91% of control (*P* < 0.0001) (**Fig. 1B**). Consistently, decreased CopGFP signals were also observed under confocal microscopy (**Fig. 1C**). Further immunoblotting analysis confirmed that MH reduced APP protein levels in a dose-dependent manner, with reductions of 22.0% and 93.6% observed at 0.25 µM and 4 µM, respectively (**Fig. 1D, E**). To examine whether inhibition of APP also affects LRRK2 levels via the APP–LRRK2 regulatory loop, we analyzed LRRK2 protein expression. Notably, MH treatment also significantly reduced LRRK2 protein levels by 74.11% (*P* =0.0077) at 1 µM (**Fig. 1D, F**). Together, these results demonstrate that MH suppresses both APP transcription and LRRK2 expression through the APP–LRRK2 regulatory loop.

**Fig. 1.**
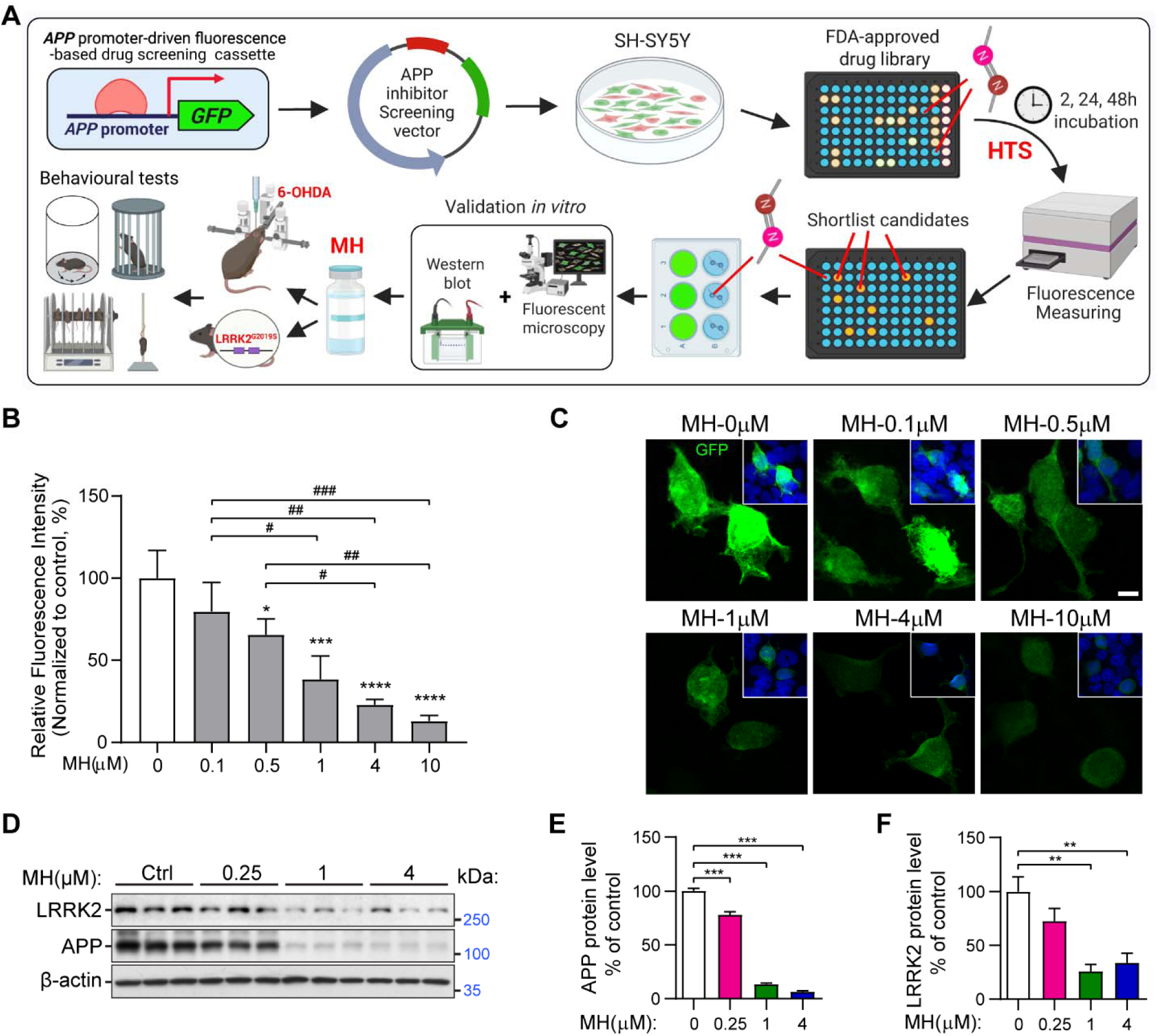
High-throughput screening identifies MH as a dual APP/LRRK2 inhibitor. **(A)** A diagram illustrates the high-throughput screening and the following validation *in vitro* and *in vivo.* (**B**) Different concentration of drug MH (0, 0.1, 0.5, 1, 4, 10 µM) was added to *APP*_pro_-CopGFP transfected SH-SY5Y cells in a 96-well plate for 24 h. GFP intensity was measured and normalized to control group. * indicates comparison to the 0 µM control group; # indicates comparison between the specified treatment groups. (**C**) *APP*_pro_-CopGFP transfected SH-SY5Y cells were treated with different concentrations of drug MH and visualized with a confocal microscopy after stained with DAPI. Scale bar = 10 µm. (**D**) MH was added to SH-SY5Y cells for 24 hours and collected for Western blot analysis. (**E-F**) Quantification of APP (**E**) and LRRK2 (**F**) from western blot in **D**. Data are presented as the mean ± SD; n=3 per group. \**P* < 0.05, \*\**P* < 0.01, and \*\*\**P* < 0.001 by one-way ANOVA with Tukey’s post *hoc* test.

### MH suppresses LRRK2 expression and phosphor-Rab10 in vitro

To further evaluate the inhibitory effects of MH on APP and LRRK2, we tested multiple cell models across different doses. In SH-SY5Y cells, the half inhibitory concentration (IC_50)_ values for endogenous APP and LRRK2 were 0.004 µM and 1.28 µM, respectively (**Fig. 2A, B**). In HEK293 cells, MH exhibited higher IC_50_ values in LRRK2^G2019S^ compared to LRRK2^WT^ transfected cells (in LRRK2^WT^ transfected cells, IC_50_ for LRRK2 and APP are 0.007 µM and 0.011 µM respectively; in LRRK2^G2019S^ transfected cells, IC_50_ for LRRK2 and APP are 3.306 µM and 0.042 µM respectively; **Fig. 2C, D**). To assess patient relevance, we treated iPSC-derived dopaminergic neurons from a PD patient carrying the LRRK2^G2019S^ mutation. MH inhibited APP and LRRK2 expression with IC_50_ values of 0.0018 µM and 0.991 µM, respectively (**Fig. 2E, F**). These findings demonstrate that MH suppresses both APP and LRRK2 expression in a dose-dependent manner across different cellular contexts, including patient-derived neurons.

**Fig. 2.**
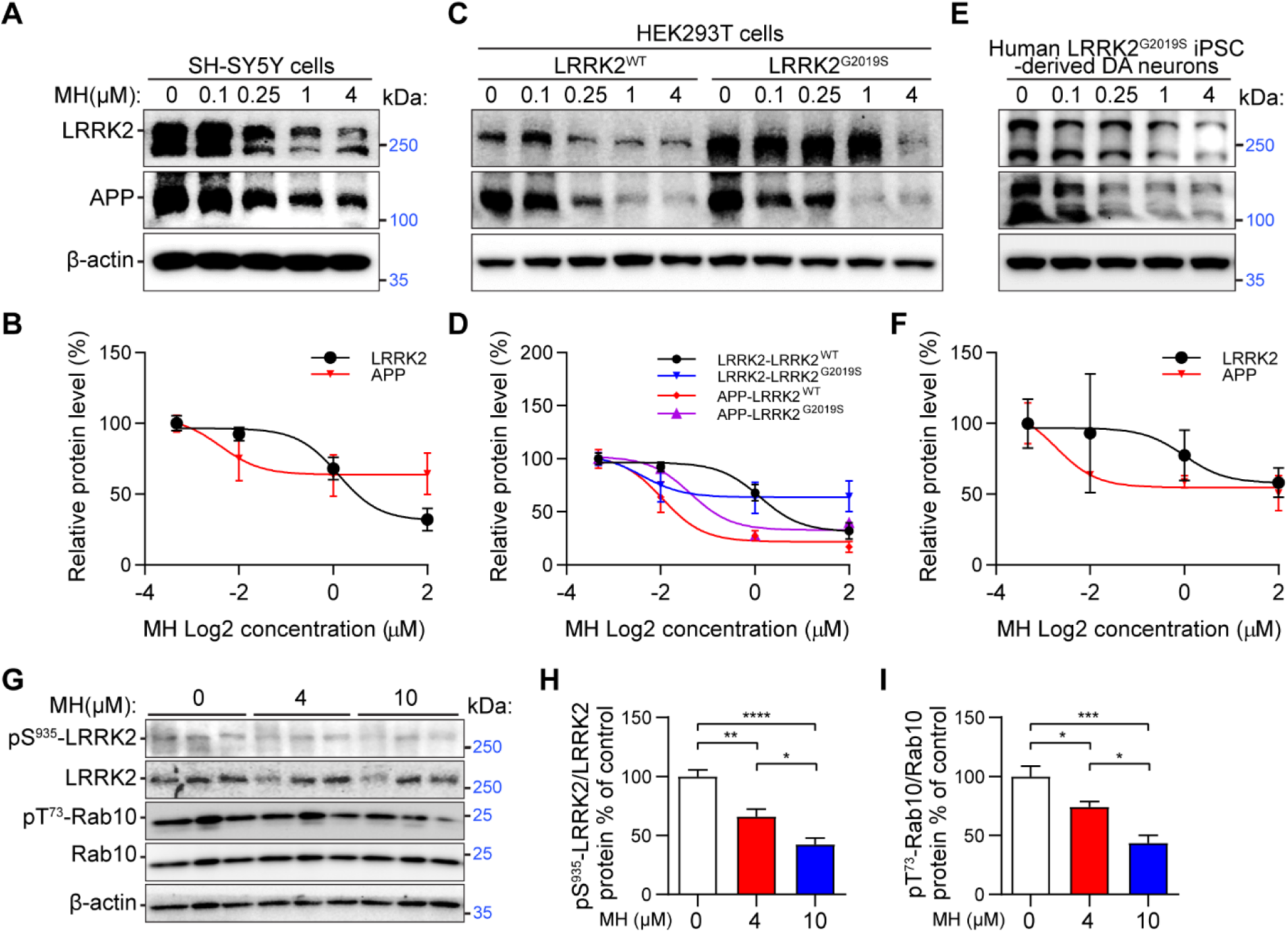
Drug MH inhibits APP/LRRK2 expression in different cells. (**A**) MH inhibits endogenous LRRK2 and APP protein levels in SH-SY5Y cells. (**B**) Relative protein amounts were plotted in Nonlin fit curve. n=3. IC_50_ for LRRK2 and APP is 1.28 and 0.004 µM respectively. (**C**) HEK293T cells were transfected with LRRK2^WT^ and LRRK2^G2019S^ plasmids and treated with 0, 0.1, 0.25, 1, and 4 µM of drug MH. After 24 h, cells were collected for immunoblotting analysis. (**D**) Relative protein amounts were plotted in Nonlin fit curve. n=3. IC_50_ for LRRK2 and APP in LRRK2^WT^ is 0.007 and 0.011 µM respectively. IC_50_ for LRRK2 and APP in LRRK2^G2019S^ is 3.306 and 0.042 µM respectively. (**E**) LRRK2^G2019S^ patient iPSC-derived DA neurons were treated with 0, 0.1, 0.25, 1, and 4 µM of drug MH. (After 24 h, cells were collected for immunoblotting analysis. (**F**) Relative protein amounts were plotted in Nonlin fit curve. n=3. IC_50_ for LRRK2 and APP in LRRK2^WT^ is 0.991 and 0.0018 µM respectively. (**G**) Healthy patient PBMCs were treated with 4 and 10 µM of drug MH for 1h before western blot. (**H-I**) Quantification of pS^935^-LRRK2 relative to LRRK2 (**H**) and pT^73^-Rab10 relative to Rab10 (**I**) from western blot in **G**. Data are presented as the mean ± SD; n=5 per group. \**P* < 0.05, \*\**P* < 0.01, \*\*\**P* < 0.001, and \*\*\*\**P* < 0.0001 by one-way ANOVA with Tukey’s post *hoc* test.

Given that MH reduces LRRK2 protein levels, we next asked whether it also affects LRRK2 kinase activity. Phosphorylation at Ser935 (pS935) is a key regulator of LRRK2 stability, 14-3-3 protein binding, and subcellular localization [34, 35]. In PD-linked LRRK2 variants, loss of pS935 is associated with dysregulated kinase activity and early pathological changes such as impaired dopamine turnover and α-synuclein accumulation [34]. In parallel, Rab10 phosphorylation at Thr73 (pT73 Rab10) serves as a direct readout of LRRK2 kinase activity and is elevated in cells and tissues carrying PD-associated mutations [19]. To assess the impact of MH, we treated human peripheral blood mononuclear cells (PBMCs) with increasing concentrations of the drug. MH reduced pS935 LRRK2 levels by 34.15% at 4 µM (*P* = 0.0036) and 57.53% at 10 µM (*P* < 0.0001; **Fig. 2G, H**), while pT73 Rab10 levels were decreased by 25.90% at 4 µM (*P* = 0.0456) and 56.25% at 10 µM (*P* = 0.002; **Fig. 2G, I**). These results demonstrate that MH effectively suppresses LRRK2 kinase activity in human cells.

Collectively, these results demonstrate that MH potently inhibits APP and LRRK2 expression across diverse cellular models, including patient-derived neurons, and effectively suppresses LRRK2 kinase activity in human cells, highlighting its therapeutic potential for targeting the APP–LRRK2 pathogenic axis in PD.

### MH inhibits LRRK2 through direct binding to its kinase domain

In addition to the inhibition of LRRK2 through the APP-LRRK2 axis, a second potential mechanism for reduced LRRK2 levels is direct inhibition of LRRK2 by MH. To test this hypothesis, we utilized computational modeling of potential binding between LRRK2 and drug MH. LRRK2 is a multidomain enzyme belonging to the leucine-rich repeat kinase family and contains several structural motifs, including the Roc GTPase domain, the kinase domain (KD), a seven-bladed WD40 repeat region, and a leucine-rich repeat (LRR) region [36]. The crystal structure of human LRRK2 (PDB ID: 8FO7) was retrieved from the Protein Data Bank and analysed to identify potential ligand-binding sites [37]. Binding site prediction using Schrödinger’s SiteMap module identified four distinct potential ligand-binding pockets distributed across the kinase, Roc, WD40, and LRR domains. These sites were evaluated based on their SiteScore and Dscore values to assess druggability [38]. MH was subsequently docked into all predicted pockets using the Glide module to determine its preferred binding location and interaction pattern [28].

Among the four identified sites, the ATP/GTP-binding pocket within the kinase domain exhibited the most favorable docking score (–7.57 kcal/mol), indicating the strongest predicted binding affinity (**Fig. 3A**). The binding mode revealed extensive hydrogen bonding and hydrophobic contacts stabilizing MH within the catalytic cleft (**Fig. 3B**). Specifically, MH formed hydrogen bonds with residues E1948, S1951, and N1999, anchoring the ligand within the catalytic cleft. The dihydroxyanthraquinone ring of MH occupied a hydrophobic cavity formed by L1885, V1893, A1904, M1947, R1957, D1994, L1996, H1998, Y2018, and T2035 which further stabilized the binding pose through hydrophobic interactions. These docking results suggest that MH preferentially binds to the kinase domain of LRRK2, likely inhibiting its catalytic activity and reinforcing its role as a key therapeutic target in PD.

**Fig. 3.**
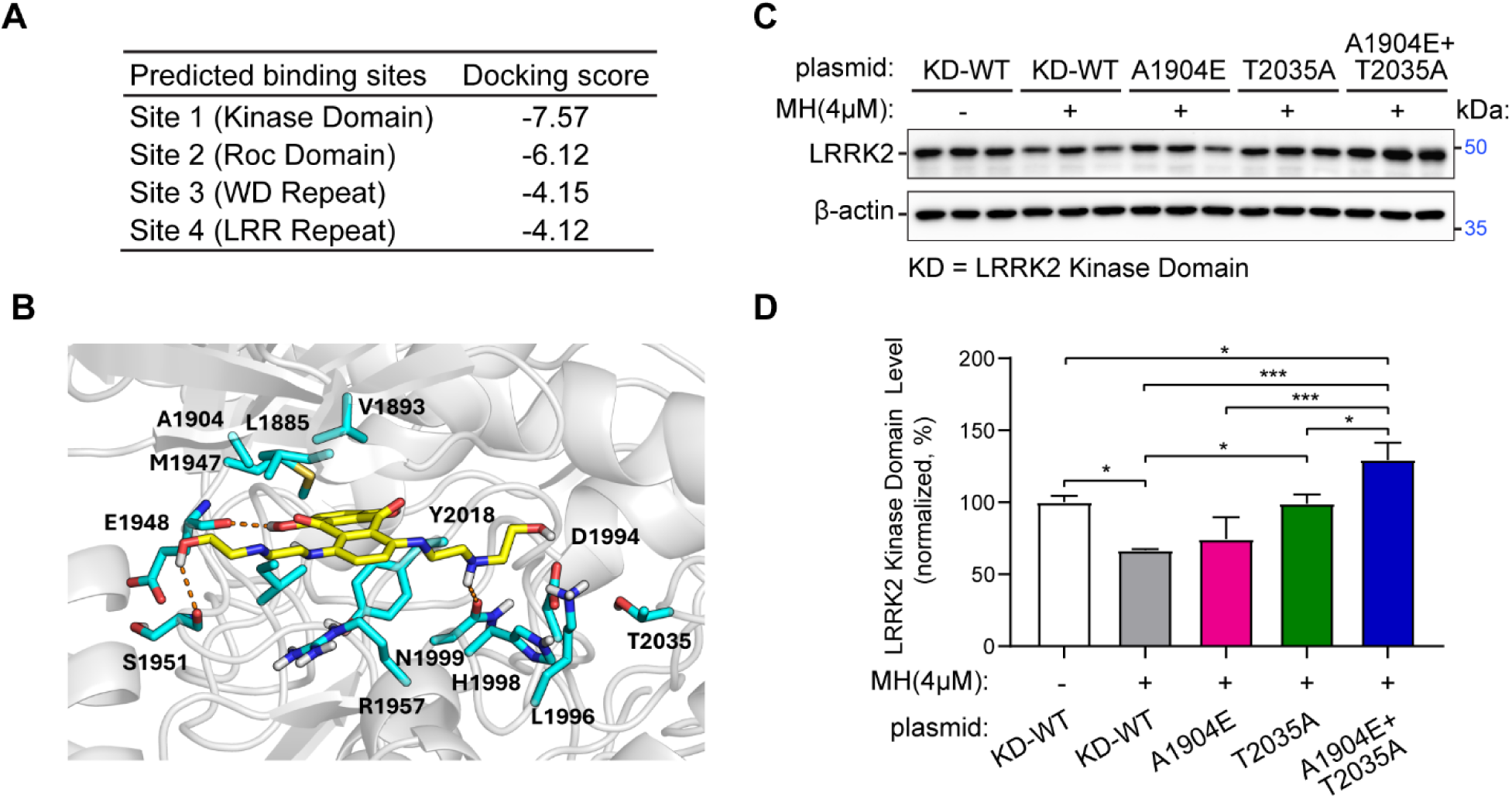
Drug MH inhibits LRRK2 through direct binding to its kinase domain. **(A)** The docking scores of MH against predicted binding sites of LRRK2. (**B**). The predicted binding pose of MH against binding site 1 (in kinase domain) of LRRK2. MH formed hydrogen bonds with E1948, S1951, and N1999 and dihydroxyanthraquinone ring of MH occupied the hydrophobic region within the predicted binding pocket. The hydrophobic region within the predicted binding pocket is formed by L1885, V1893, A1904, M1947, R1957, D1994, L1996, H1998, Y2018, and T2035 residues. (**C**) A1904E and T2035A were mutated in a LRRK2 kinase domain (KD)-expressing plasmid by site-directed mutagenesis. SH-SY5Y cells were transfected with LRRK2 KD and KD with mutations. After 24 h of 4 µM MH treatment, cells were collected for western blot analysis. (**D**) Quantification of LRRK2-KD from western blot in **C**. Data are presented as the mean ± SD; n=3 per group. \**P* < 0.05 and \*\*\**P* < 0.001 by one-way ANOVA with Tukey’s post *hoc* test.

To further predict the importance of individual amino acid residues for ligand binding, mutational analysis was performed using the Residue Scanning tool in Schrödinger. Residues within 5.0 Å of the ligand were systematically mutated, and changes in binding affinity (ΔΔG) were calculated using MM-GBSA refinement for both wild-type and mutant complexes. The analysis identified A1904, R1957, H1998, N1999, and T2035 as key contributors to ligand binding, providing mechanistic insight into the stabilization of MH within the kinase domain. To validate this interaction, we introduced point mutations A1904E and T2035A, which exhibited the highest MM-GBSA scores in the simulation, into the LRRK2 kinase domain using site-directed mutagenesis to disrupt ligand binding. The T2035A mutation significantly increased LRRK2 KD protein levels, fully restoring them to the LRRK2 KD WT baseline without MH treatment (**Fig. 3C, D**). The double mutation (A1904E/T2035A) further elevated protein levels compared to either single mutation alone (**Fig. 3C, D**). These findings suggest that Mitoxantrone stabilizes within the catalytic pocket through a combination of polar and hydrophobic interactions, consistent with its potential role as a kinase inhibitor.

Collectively, our findings indicate that MH attenuates neurotoxicity both by modulating the APP–LRRK2 axis and through direct LRRK2 inhibition, effectively disrupting the APP–LRRK2 toxic loop via dual inhibition of APP and LRRK2.

### MH rescues DA neuronal loss and motor deficits in PD mice

Since MH inhibits LRRK2 protein levels, kinase activity, and downstream targets, we next investigated its effects in PD mouse models. 6-Hydroxydopamine (6-OHDA) is a neurotoxin commonly used to model PD by selectively destroying dopaminergic neurons in the nigrostriatal pathway, producing motor and biochemical deficits reminiscent of PD pathology [39]. Our previous work showed that inhibiting LRRK2 via the APP-LRRK2 loop rescued DA neuronal loss and motor deficits in 6-OHDA-induced PD mice [21]. Clinically, MH is administered at 12–14 mg/m² intravenous infusion (IV) every 21 days for acute non-lymphocytic leukemia, 12 mg/m² IV every 21 days for hormone-refractory prostate cancer, and 12 mg/m² every 3 months (maximum cumulative dose ∼140 mg/m²) for secondary progressive or relapsing multiple sclerosis [40–42]. To generate the PD model, 4 µg of 6-OHDA was stereotaxically injected into the left striatum. After confirming the PD phenotype by rotation test, mice received intraperitoneal MH at 1 mg/kg per day for 5 days (cumulative dose 5 mg/kg), corresponding to a human dose of about 15 mg/m² [24, 43]. Two weeks post-treatment, behavioral assessments and immunoblotting analyses were conducted (**Fig. 4A**). MH treatment restored tyrosine hydroxylase (TH) levels in the left striatum to those of the intact right striatum, indicating recovery of DA neurons (**Fig. 4B, C**). In the cylinder test, the left rearing ratio in MH-treated mice decreased to 58.91% (P = 0.012) versus controls (**Fig. 4D**), and treated mice descended the pole more quickly (**Fig. 4E**), reflecting improved motor function.

**Fig. 4.**
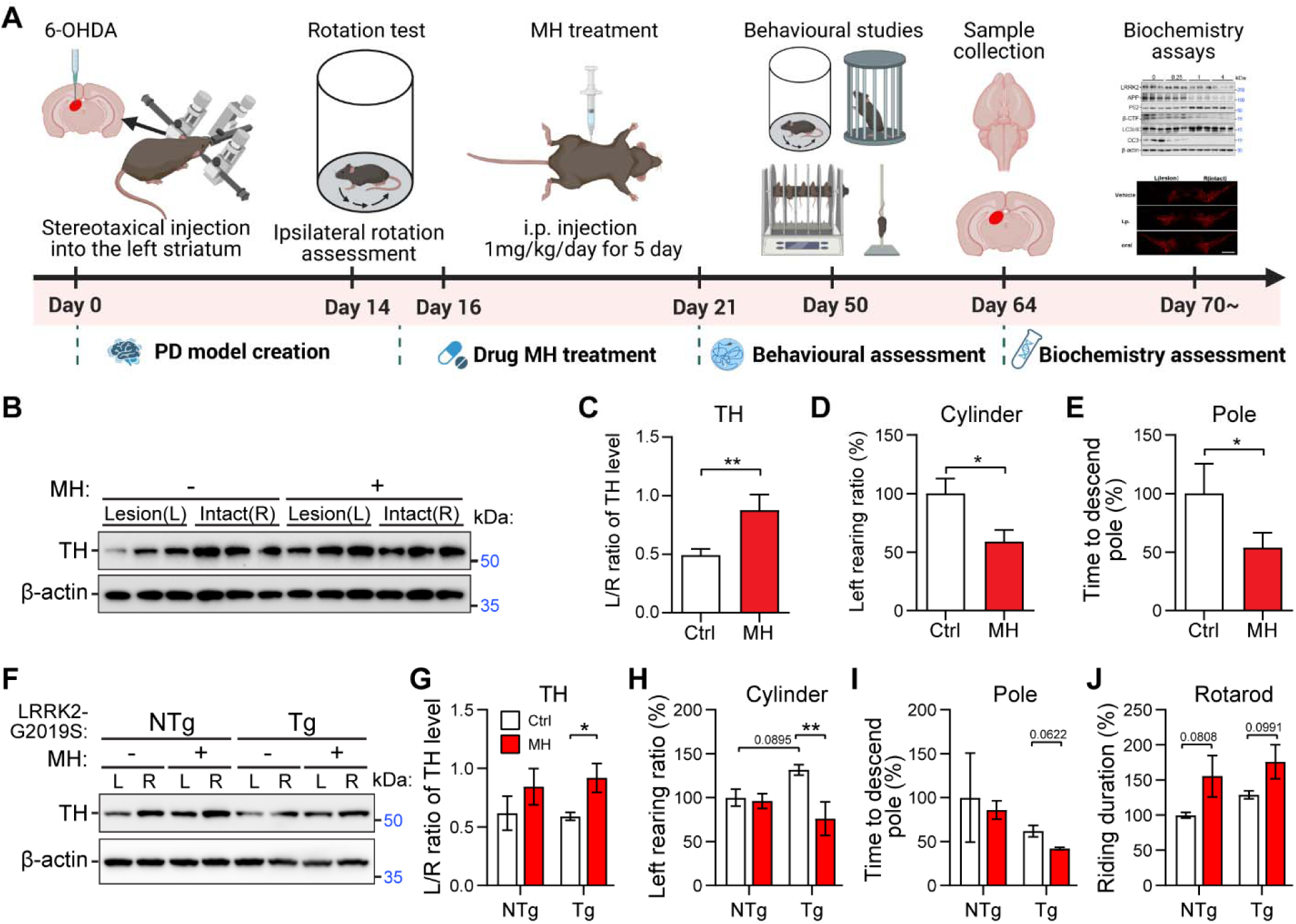
Drug MH ameliorates neurotoxicity and behavioral deficits in PD mice. (**A**) A diagram illustrates the procedures of drug MH treatment on 6-OHDA PD mice. (**B-E**) 6-OHDA (4 μg) was stereotaxically injected into 4-month-old C57 mice on the left striatum (day 0), and MH (1 mg/kg) was administered via i.p. on day 8 for 5 days of treatments before Cylinder test and Pole test. Brain and tissues were then harvested for western blot. Western blotting images of relative TH expression in left and right striata. (**C**) Quantification of TH from western blot in **B**. Behavioural tests were performed two weeks post drug treatment. Left to total rearing in Cylinder test (**D**) and time to descend pole in the Pole test (**E**) were plotted. Data are mean ± SD; n=3 per group. \**P* < 0.05 and \*\**P* < 0.01 by two-tailed Student’s *t* test. (**F-J**) 6-OHDA (2 μg) was stereotaxically injected into 4-month-old LRRK2^G2019S^ on the left striatum (day 0), and MH (1 mg/kg) was administered via i.p. on day 8 for 5 days of treatments before behavioral tests. Brain and tissues were then harvested for western blot (**F**). (**G**) Quantification of TH from western blot in **F**. Behavioural tests were performed two weeks post drug treatment. Left to total rearing in Cylinder test (**H**), time to descend pole in the Pole test (**I**), and riding duration in the Rotarod test (**J**) were plotted. Data are mean ± SD; n=3 per group. \**P* < 0.05 and \*\**P* < 0.01 by two-way ANOVA with multiple comparison.

To examine effects in a genetic PD model, MH was administered to LRRK2^G2019S^ transgenic (Tg) mice, which exhibit subtle early PD phenotypes without overt dopaminergic degeneration [44–46]. To accelerate pathology, half-dose 6-OHDA (2 µg) was injected. MH treatment increased striatal TH levels relative to non-treated LRRK2^G2019S^ Tg mice (**Fig. 4F, G**) and reduced left rearing ratio in the cylinder test (**Fig. 4H**). Treated mice also showed trends toward improved motor coordination in the pole and rotarod tests (**Fig. 4I, J**).

Overall, these results indicate that MH rescues DA neuronal loss and ameliorates motor deficits in both 6-OHDA-induced and LRRK2^G2019S^ PD mouse models, supporting its potential as a therapeutic for both sporadic and LRRK2-linked familial PD.

### Oral MH protects against dopaminergic neuron loss and rescues motor deficits in PD mice

MH is FDA-approved for intravenous infusion in cancer and multiple sclerosis, but its clinical use is constrained by dose-dependent cardiotoxicity in some patients [47–49]. To mitigate potential adverse effects and enhance translational relevance, we reduced the i.p. injection dose by half and tested oral administration, which is more patient-friendly, non-invasive, cost-effective, and suitable for long-term treatment [50, 51]. Behavioral assessments were conducted at 2, 6, and 12 weeks after MH treatment. Both cumulative i.p. injection of 2.5 mg/kg (0.5 mg/kg daily for 5 days) and oral gavage of 5 mg/kg (1 mg/kg daily for 5 days) significantly attenuated ipsilateral rotations in 6-OHDA mice across all time points, with the strongest effect at 12 weeks (61.24%, *P* < 0.001 for i.p.-2.5 and 71.95%, *P* < 0.0001 for oral-5 vs. vehicle; **Fig. 5A**). Similar trends were observed in the cylinder test, where maximal inhibition was reached at 6 weeks (78.45%, *P* < 0.001 and 79.17%, *P* < 0.001, respectively; **Fig. 5B**). In the rotarod test, oral MH significantly improved motor performance as early as 2 weeks, while i.p.-2.5 showed benefits only after 6 weeks (**Fig. 5C**). Likewise, oral MH produced significant improvements in the pole test at 6 and 12 weeks, whereas i.p.-2.5 showed only a modest trend (**Fig. 5D**). These results suggest that both regimens improved motor performance, with oral MH showing superior efficacy.

**Fig. 5.**
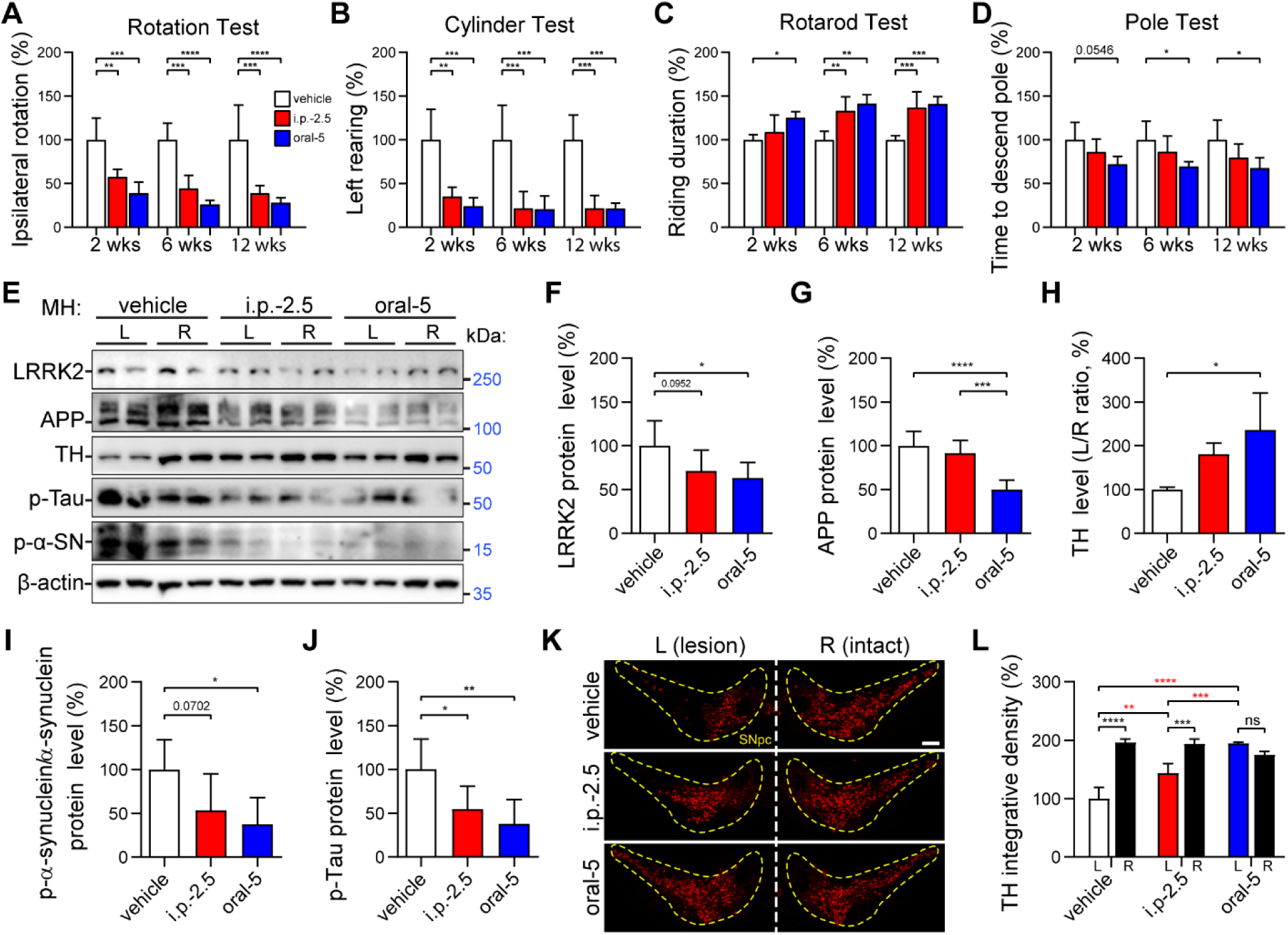
Oral administration of MH ameliorates neurotoxicity and motor behavioral deficits in PD mice. 6-OHDA (4 μg) was stereotaxically injected into 4-month-old C57 mice on the left striatum (day 0), and MH was administered via i.p. (0.5 mg/kg) or oral gavage (1 mg/kg) on day 8 for 5 days (cumulative doses of i.p. 2.5 mg/kg and oral 5 mg/kg respectively) before behavioral test after 2-, 6-, and 12-weeks of MH treatment. (**A-D**). 2-, 6-, and 12-weeks after MH treatment, anti-clockwise ipsilateral rotation in the Rotation Test (**A**), left rearing ratio in the Cylinder Test (**B**), riding duration in the Rotarod Test (**C**), and time to descend the pole in the Pole Test (**D**) were plotted. For each time point, animal behavioural was normalized to the vehicle control. Data are presented as the mean ± SD; n=4 per group. \**P* < 0.05, \*\**P* < 0.01, \*\*\**P* < 0.001, and \*\*\*\**P* < 0.0001 by one-way ANOVA with Tukey’s post *hoc* test. (**E**) After performing behavioural tests 12-weeks post drug treatment, mice were sacrificed. The striatum tissues from left side lesion (L) and intact right side (R) of the mouse brain were used for western blot analysis. (**F-J**) Quantification of LRRK2 (**F**), APP (**G**), TH (**H**), pS^129^-α-synuclein to α-synuclein (**I**), and pS^199/202^-Tau (**J**) from western blot in **E**. Protein levels are presented as the left to right ratio and normalized to the vehicle group. Data are presented as the mean ± SD; n=3-6 per group. \**P* < 0.05, \*\**P* < 0.01, \*\*\**P* < 0.001, and \*\*\*\**P* < 0.0001 by one-way ANOVA with Tukey’s post *hoc* test. (**K**) After performing behavioural tests 12-weeks post drug treatment, mice were sacrificed. The mouse brains were immunohistochemical stained with TH. The dash line area represents the SNpc region. Scale bar = 100 μm. (**L**) Quantification of integrative TH+ DA neuron density in **K**. Data are presented as the mean ± SD; n=3. \*\**P* < 0.01, \*\*\**P* < 0.001, and \*\*\*\**P* < 0.0001 by two-way ANOVA with multiple comparison.

We next evaluated neuropathological changes at 12 weeks post-treatment. Oral MH robustly reduced striatal LRRK2 and APP levels, whereas i.p.-2.5 had only a partial effect (**Fig. 5E–G**). TH expression was fully restored after oral treatment (**Fig. 5E, H**). Importantly, phospho-α-synuclein was significantly reduced by oral MH but only modestly by i.p.-2.5 (**Fig. 5E, I**). Both regimens decreased phospho-Tau in the striatum (**Fig. 5E, J**). In the SNpc, the number of TH□ DA neurons on the lesioned side increased 1.46-fold following i.p. 2.5 mg/kg treatment compared to vehicle (**Fig. 5K, L**). Notably, oral gavage of 5 mg/kg fully restored DA neuron numbers to levels comparable to the intact side and yielded a 2.18-fold increase over the vehicle group (**Fig. 5K, L**).

Collectively, these findings demonstrate that halved i.p. dosing of MH still confers behavioral and neuroprotective benefits in 6-OHDA mice, while oral administration at 5 mg/kg provides superior recovery of motor function and dopaminergic integrity, along with reductions in phospho-α-synuclein and phospho-Tau. This highlights the potential of oral MH as a promising therapeutic strategy for PD and possibly AD.

### Oral MH is well tolerated with no adverse effects on vital organs or intestinal integrity

MH is an antineoplastic agent originally developed for cancer treatment and later approved for secondary progressive and relapsing-remitting multiple sclerosis [52, 53]. Its principal adverse effect is dose-dependent cardiotoxicity [54, 55]; accordingly, clinical use is limited by a maximum cumulative lifetime dose of 140 mg/m² [56]. In this study, MH was administered at i.p. 2.5–5 mg/kg and oral 5 mg/kg, and we assessed potential toxicity in major organs (heart, brain, kidney, liver, thymus, spleen, lung) as well as possible gastrointestinal effects from oral dosing. After 12 weeks, we measured body weight and the organ-to-body weight ratio (organ index), a standard toxicology metric for detecting drug-induced organ changes [57, 58]. Body weight was unchanged in both i.p.-2.5 and oral-5 groups over the 3-month follow-up (**Fig. 6A**). Oral MH (5 mg/kg) did not alter any organ indices (**Fig. 6B–H**), whereas i.p.-2.5 produced a slight increase in heart-to-body weight ratio compared with vehicle (**Fig. 6H**), which may reflect early cardiac adaptation or stress consistent with known MH cardiotoxicity [54, 59].

**Fig. 6.**
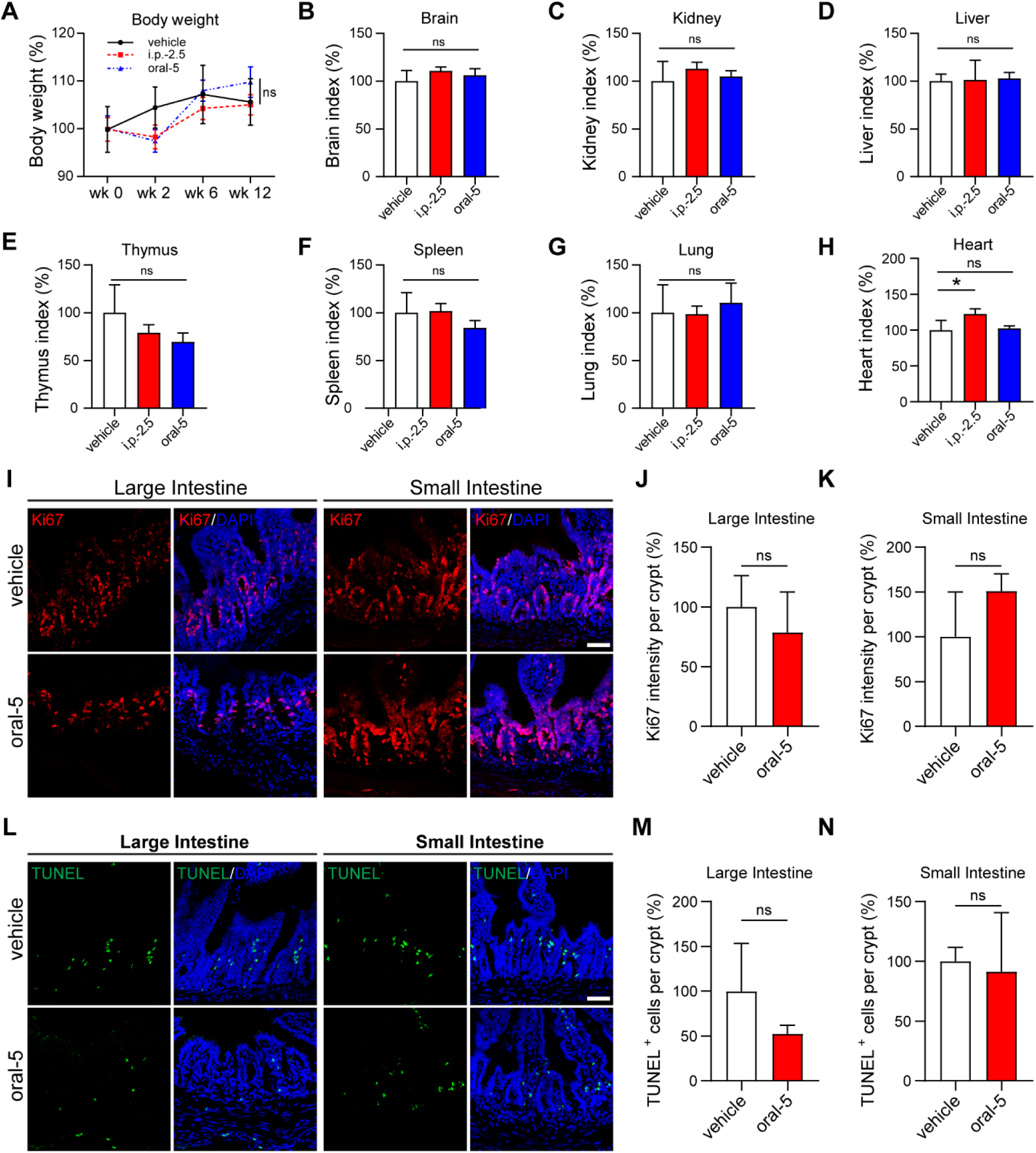
Oral MH is well tolerated with no adverse effects on vital organs or intestinal integrity. **(A**) Body weight was unchanged in i.p.-2.5 or oral-5 mg/kg groups during 2-, 6-, and 12-weeks of treatment. (**B–H**) After 12-weeks of drug treatment, weights of vital organs and whole body of mice were measured. Organ indices (organ-to-body weight ratios) of brain (**B**), kidney (**C**), liver (**D**), thymus (**E**), spleen (**F**), lung (**G**), and heart (**H**). Data are mean ± SD; n=4 mice per group. \**P* < 0.05 by one-way ANOVA with Tukey’s post hoc test. ns = not significant. (**I**) After 12-weeks of drug treatment, tissues from large intestine and small intestine were cryosectioned immunohistochemical staining. (**J-K**) Quantification of Ki67 intensity per crypt in large intestine (**J**) and small intestine (**K**) in **I**. (**L**) TUNEL assays in large intestine and small intestine. (**M-N**) Quantification of TUMEL^+^ cells per crypt in large intestine (**M**) and small intestine (**N**) in **L**. Data are presented as mean ± SD; ns = not significant by two-tailed student’s *t* test. Scale bar = 50 µm.

To evaluate potential gastrointestinal toxicity after oral MH, we performed Ki67 staining to assess epithelial proliferation and TUNEL assays to quantify apoptosis in monolayer sections of small and large intestine [60]. Ki67 intensity per crypt was similar between oral-5 and vehicle groups in both intestinal segments (**Fig. 6I–K**). Likewise, there was no significant difference in TUNEL^+^ cell counts per crypt between groups (**Fig. 6L–N**).

Overall, these data indicate that oral MH at 5 mg/kg produced no detectable toxicity in major organs or the intestinal epithelium, while the modest increase in heart index after low-dose i.p. administration warrants further functional cardiac assessment given MH’s known cardiotoxic potential.

### Oral MH shows no detectable cardiotoxicity in PD mice

MH has been reported to cause dose-dependent cardiotoxicity, including congestive heart failure and reduced left ventricular ejection fraction in human clinical trials [55]. In mice, cumulative i.p. administration of 7 mg/kg MH has been associated with cardiac toxicity [54]. To further assess potential cardiac side effects at lower doses, we performed histopathological evaluations of heart tissue. No pathological alterations in myocardial morphology or cardiomyocyte structure were observed among the i.p. 2.5 mg/kg, oral 5 mg/kg, and vehicle-treated groups (**Fig. 7A**). Consistently, Sirius Red staining showed no significant differences in the collagen-to-skeletal muscle area ratio, indicating that neither i.p. 2.5 mg/kg nor oral 5 mg/kg MH induced myocardial fibrosis (**Fig. 7B, C**). To investigate whether MH triggered cardiac inflammation, we performed CD68 immunostaining to detect pro-inflammatory M1 macrophages [61, 62]. CD68□ area was comparable between oral 5 mg/kg MH and vehicle groups, whereas i.p. 2.5 mg/kg MH showed a modest downward trend in CD68□ staining (**Fig. 7D, E**). This reduction is consistent with MH’s well-established immunosuppressive and anti-inflammatory activity, including its clinical use in multiple sclerosis to suppress immune cell infiltration and inflammation [63, 64].

**Fig. 7.**
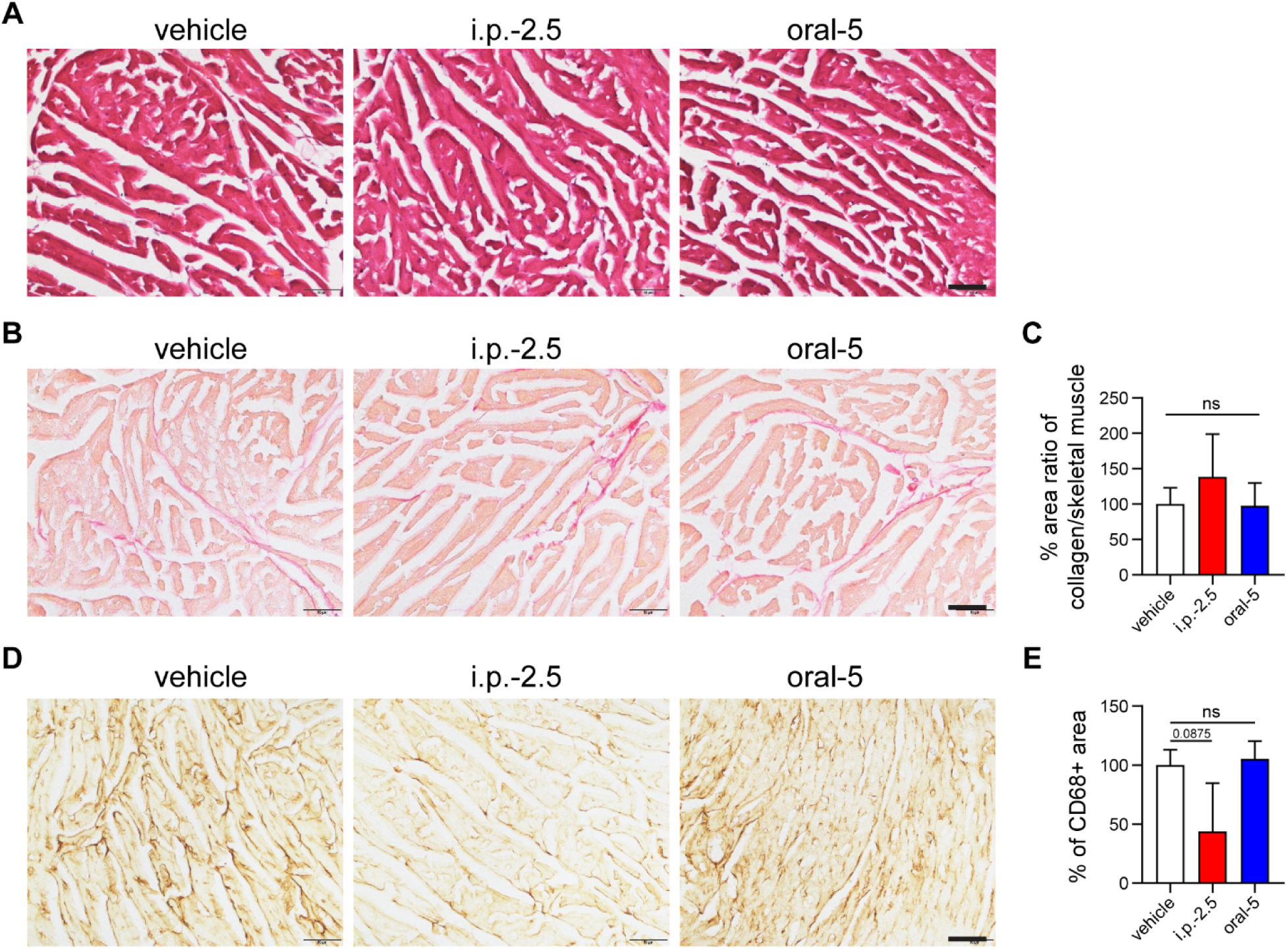
Oral MH shows no detectable cardiotoxicity in PD mice. (**A**) Representative hematoxylin and eosin (H&E) staining images of heart sections showed no pathological changes in morphology or cardiomyocytes among i.p. 2.5 mg/kg, oral 5 mg/kg, and vehicle groups. (**B–C**) Sirius Red staining demonstrated no significant differences in collagen-to-skeletal muscle area ratio among treatment groups, indicating the absence of myocardial fibrosis. (**D–E**) Immunostaining for CD68+ macrophages revealed comparable inflammatory cell infiltration between oral 5 mg/kg MH and vehicle groups, while i.p. 2.5 mg/kg MH showed a slight reduction in CD68+ area. Data are presented as mean ± SD; n=3 per group. \**P* < 0.05 by one-way ANOVA with Tukey’s post hoc test. ns = not significant. Scale bar = 50 µm.

Taken together, these findings suggest that both oral 5 mg/kg and i.p. 2.5 mg/kg MH do not induce cardiac toxicity under the tested conditions. Moreover, oral administration appears to exert minimal adverse effects on the heart, further supporting its safety advantage.

## DISCUSSION

In this study, we identify the FDA-approved drug MH as a potent, dual inhibitor of the APP-LRRK2 pathogenic axis, demonstrating its significant neuroprotective and behavioral benefits in preclinical models of Parkinson’s disease. Our findings build upon our previous work that established a self-reinforcing cycle between APP/AICD and LRRK2 as a key driver of PD pathology. By employing a high-throughput screening strategy, we discovered that MH simultaneously suppresses APP transcription and LRRK2 protein levels, effectively disrupting this feed-forward loop at its origin [20, 21]. Furthermore, we elucidated a second, direct mechanism whereby MH binds to the kinase domain of LRRK2, inhibiting its enzymatic activity. This dual-action mechanism—simultaneously targeting the expression of both proteins and the kinase function of LRRK2—represents a comprehensive therapeutic strategy that may be more efficacious than targeting either component alone.

The therapeutic potential of MH was consistently demonstrated across multiple experimental setups. In cellular models, including patient-derived iPSC dopaminergic neurons, MH not only reduced APP and LRRK2 but also mitigated downstream neurotoxic events, including aberrant autophagy and apoptosis. Furthermore, MH treatment in human PBMCs significantly reduced phosphorylation of LRRK2 at Ser935 and its substrate Rab10 at Thr73, establishing its efficacy in suppressing LRRK2 kinase activity, a key driver of pathology [19]. The rescue of dopaminergic neurons, as evidenced by restored tyrosine hydroxylase levels, and the amelioration of motor deficits in both 6-OHDA-induced and LRRK2^G2019S^ genetic mouse models substantiate its disease-modifying potential. Importantly, we established that oral administration of MH, a more practical and patient-friendly route, conferred superior neuroprotective and functional recovery compared to intraperitoneal injection. This was accompanied by a significant reduction in key pathological proteins, including phospho-α-synuclein and phospho-Tau [31], suggesting that MH’s benefits may extend beyond the dopaminergic system and could be relevant for other neurodegenerative disorders.

PD and AD share overlapping neuropathological features. Dementia develops in approximately 25–30% of PD patients, and Parkinson’s disease dementia (PDD) accounts for about 3.6% of all dementia cases among the elderly population [65–67]. Moreover, neuropathological analyses have revealed Lewy body pathology—the hallmark of PD—in 40–60% of AD cases, particularly within limbic and entorhinal regions, although neocortical involvement occurs less frequently [68]. Our previous studies identified the AICD/APP–LRRK2 regulatory axis as a convergent pathogenic pathway implicated in both AD and PD [20, 21]. Targeting this axis offers therapeutic potential across multiple neurodegenerative conditions. We previously demonstrated that itanapraced (also called CHF5074), an AICD inhibitor that blocks AICD nuclear translocation and ameliorates neuroinflammation and memory deficits in transgenic AD models and patients [22], also modulates the AICD/APP–LRRK2 loop and attenuates neurodegeneration in PD models [21]. In the present study, we further provide evidence that targeting this shared regulatory pathway alleviates neuronal loss and motor deficits in PD. Given that MH treatment also reduces APP and phosphorylated Tau—key pathological markers of AD—it is plausible that MH may exert similar therapeutic benefits in AD. Supporting this notion, a previous study reported that mitoxantrone inhibits Aβ seeding-mediated aggregation and stabilizes diffuse plaques in AD mouse models [69]. Together, these findings suggest that MH represents a promising repurposed therapeutic candidate for both AD and PD, as well as other APP- and LRRK2-associated neurodegenerative disorders.

A critically important finding of our work is the demonstration that oral administration of MH is not only feasible but also highly effective, and well-tolerated. Oral delivery conferred superior neuroprotective and motor benefits compared to intraperitoneal injection, a key consideration for patient compliance and long-term chronic therapy [50, 51]. Comprehensive toxicological assessments revealed no significant damage to vital organs or the gastrointestinal tract at the efficacious oral dose of 5 mg/kg. Most notably, and in contrast to the known dose-dependent cardiotoxicity of MH in its oncological use [54, 55], we observed no histopathological evidence of myocardial fibrosis or inflammation at this regimen, suggesting a favorable safety window for its potential repurposing in neurodegenerative diseases.

A critical consideration for repurposing MH is its known risk of dose-dependent cardiotoxicity [47, 70]. Our investigation into the safety profile of the lower doses used in this study is therefore highly encouraging. We found that oral administration at 5 mg/kg did not induce detectable cardiotoxicity, organ damage, or gastrointestinal injury over a 12-week period. Several preclinical and clinical studies indicate that lower cumulative doses or modified delivery of MH may avoid overt myocardial fibrosis or inflammatory infiltration. Mice treated at modest cumulative doses show only mild inflammatory marker elevation without heavy infiltration or fibrosis [54], and multiple sclerosis patients receiving 12 mg/m² have generally preserved cardiac function with rare declines in left ventricular ejection fraction (LVEF) in the first year of treatment [71]. The absence of myocardial fibrosis and inflammatory infiltration at these doses suggests a potential therapeutic window where the neuroprotective benefits of MH can be harnessed without its primary adverse effect. This positions MH favorably against its historical profile and underscores the importance of dose optimization for new indications.

Despite these promising results, our study has several limitations. The observation period, while sufficient to demonstrate efficacy, was relatively short for a chronic neurodegenerative disorder; longer-term studies are necessary to confirm sustained benefits and chronic safety. Furthermore, while our mouse models recapitulate key features of PD, they do not fully capture the slow, progressive nature of the human disease, which is an inherited drawback of mouse models [72, 73]. The precise structural determinants of the MH-LRRK2 interaction, while predicted by docking studies, would benefit from validation through crystallography. While our docking studies suggest that MH binds to specific residues in the kinase domain of LRRK2, high-resolution structural data, including cryo-EM or X-ray crystallography as reported for LRRK2-inhibitor complexes [37, 74], would be required to precisely define the binding mode and confirm the identified hydrogen bonds and hydrophobic pocket occupancy. Finally, the potential impact of MH’s known immunosuppressive effects on neuroinflammation in PD was not fully explored here and warrants further investigation.

## CONCLUSIONS

In conclusion, our work repositions MH from a chemotherapeutic agent to a promising neurotherapeutic, offering a novel and mechanistically grounded strategy to disrupt a core pathogenic pathway in PD.

## DECLARATIONS

## Acknowledgments

The graphic illustrations were created in https://BioRender.com.

## Funding

This research was supported by research grants administered by the Singapore Ministry of Health’s National Medical Research Council, including:

Open Fund-Large Collaborative Grant (LCG002–SPARK II, to EKT)

Singapore Parkinson’s Disease Programme, or Sparkle, to EKT)

Clinician Scientist Individual Research Grant (MOH-CIRG21nov-0001, to ZDZ)

Open Fund-Individual Research Grant (OFIRG23jul-0075, to ZDZ)

Open Fund-Individual Research Grant (OFIRG24jul-0093, to LZ)

## Author contributions

Conceptualization: HT, LZ

Methodology: HT, LZ, ZWZ, LZ, ZDZ, HF, YXC, PZ

Investigation: HT, ZWZ, CKJ, MYGM, WTS

Visualization: HT, CKJ

Funding acquisition: ZDZ, EKT, LZ

Project administration: SYC

Supervision: LZ, EKT

Writing: HT, CKJ

## Ethics approval and consent to participate

All protocols and procedures of the animal experiments in this study were reviewed and approved by the Institutional Review Board and the Institutional Animal Care and Use Committee (A25006) of the National Neuroscience Institute and Lee Kong Chian School of Medicine, Nanyang Technology University.

## Consent for publication

All authors read and approved the final manuscript.

## Competing interests

Authors declare that they have no competing interests.

## Data and materials availability

The authors declare that all data supporting the findings of this study are available in this article. Further inquiries can be directed to the corresponding authors.

## Abbreviations

6-OHDA: 6-Hydroxydopamine
AD: Alzheimer’s disease
AICD: APP intracellular domain
APP: Amyloid precursor protein
αSyn: alpha-synuclein
DA: dopaminergic
DMEM: Dulbecco’s Modified Eagle Medium
HTS: High-throughput screening
IHC: Immunohistochemistry
IC_50_: Half inhibitory concentration
IP: intrapetritoneal
iPSC: induced pluripotent stem cell
IV: Intravenous
KD: Kinase domain
LRRK2: Leucine-rich repeat kinase 2
MH: Mitoxantrone hydrochloride
PBMC: Peripheral blood mononuclear cell
PD: Parkinson’s disease
SN: Substantia nigra
SNpc: Substantia nigra pars compacta
TG: Transgenic
TH: Tyrosine hydroxylase

## Notes

### Competing Interest Statement

The authors have declared no competing interest.

### Summary of Updates

Removed previous Figure 3 and related result section; Updated previous Fig. 4B (now Fig. 3B); Updated methods section; Revised some of the subtitles.

## REFERENCES

1. Skibinski G, Finkbeiner S: Drug discovery in Parkinson’s disease-Update and developments in the use of cellular models. Int J High Throughput Screen 2011, 2011:15–25. 10.2147/IJHTS.S8681

2. Charvin D, Medori R, Hauser RA, Rascol O: Therapeutic strategies for Parkinson disease: beyond dopaminergic drugs. Nature Reviews Drug Discovery 2018, 17:804–822. 10.1038/nrd.2018.136

3. Wakabayashi K, Mori F, Takahashi H: Progression patterns of neuronal loss and Lewy body pathology in the substantia nigra in Parkinson’s disease. Parkinsonism & Related Disorders 2006, 12:S92–S98. 10.1016/j.parkreldis.2006.05.028

4. Brakedal B, Toker L, Haugarvoll K, Tzoulis C: A nationwide study of the incidence, prevalence and mortality of Parkinson’s disease in the Norwegian population. NPJ Parkinsons Dis 2022, 8:19. 10.1038/s41531-022-00280-4

5. Kumari U, Tan EK: LRRK2 in Parkinson’s disease: genetic and clinical studies from patients. FEBS J 2009, 276:6455–6463. 10.1111/j.1742-4658.2009.07344.x

6. Dorsey E, Constantinescu R, Thompson J, Biglan K, Holloway R, Kieburtz K, Marshall F, Ravina B, Schifitto G, Siderowf A: Projected number of people with Parkinson disease in the most populous nations, 2005 through 2030. Neurology 2007, 68:384–386. 10.1212/01.wnl.0000247740.47667.03

7. 2020 Alzheimer’s disease facts and figures. Alzheimers Dement 2020. 10.1002/alz.12068

8. Chang KH, Chen CM: The Role of Oxidative Stress in Parkinson’s Disease. Antioxidants (Basel) 2020, 9. 10.3390/antiox9070597

9. Kim TY, Lee BD: Current therapeutic strategies in Parkinson’s disease: Future perspectives. Mol Cells 2025, 48:100274. 10.1016/j.mocell.2025.100274

10. Sackner-Bernstein J: Rethinking Parkinson’s disease: could dopamine reduction therapy have clinical utility? J Neurol 2024, 271:5687–5695. 10.1007/s00415-024-12526-7

11. Kang UB, Marto JA: Leucine-rich repeat kinase 2 and Parkinson’s disease. Proteomics 2017, 17. 10.1002/pmic.201600092

12. Di Maio R, Hoffman EK, Rocha EM, Keeney MT, Sanders LH, De Miranda BR, Zharikov A, Van Laar A, Stepan AF, Lanz TA, et al: LRRK2 activation in idiopathic Parkinson’s disease. Sci Transl Med 2018, 10. 10.1126/scitranslmed.aar5429

13. Jennings D, Huntwork-Rodriguez S, Henry AG, Sasaki JC, Meisner R, Diaz D, Solanoy H, Wang X, Negrou E, Bondar VV, et al: Preclinical and clinical evaluation of the LRRK2 inhibitor DNL201 for Parkinson’s disease. Sci Transl Med 2022, 14:eabj2658. 10.1126/scitranslmed.abj2658

14. Jennings D, Huntwork-Rodriguez S, Vissers M, Daryani VM, Diaz D, Goo MS, Chen JJ, Maciuca R, Fraser K, Mabrouk OS, et al: LRRK2 Inhibition by BIIB122 in Healthy Participants and Patients with Parkinson’s Disease. Mov Disord 2023, 38:386–398. 10.1002/mds.29297

15. Biogen, Denali T: A Study to Evaluate the Efficacy and Safety of BIIB122 in Participants With Early Parkinson’s Disease (LUMA). ClinicalTrialsgov 2022.

16. Biogen, Denali T: A Study to Evaluate the Safety and Pharmacodynamic Effects of BIIB122 in Participants With LRRK2-Associated Parkinson’s Disease (BEACON). ClinicalTrialsgov 2024.

17. Alessi DR, Sammler E: LRRK2 kinase in Parkinson’s disease. Science 2018, 360:36–37. 10.1126/science.aar5683

18. Daher JP, Abdelmotilib HA, Hu X, Volpicelli-Daley LA, Moehle MS, Fraser KB, Needle E, Chen Y, Steyn SJ, Galatsis P, et al: Leucine-rich Repeat Kinase 2 (LRRK2) Pharmacological Inhibition Abates alpha-Synuclein Gene-induced Neurodegeneration. J Biol Chem 2015, 290:19433–19444. 10.1074/jbc.M115.660001

19. Steger M, Tonelli F, Ito G, Davies P, Trost M, Vetter M, Wachter S, Lorentzen E, Duddy G, Wilson S, et al: Phosphoproteomics reveals that Parkinson’s disease kinase LRRK2 regulates a subset of Rab GTPases. Elife 2016, 5. 10.7554/eLife.12813

20. Chen ZC, Zhang W, Chua LL, Chai C, Li R, Lin L, Cao Z, Angeles DC, Stanton LW, Peng JH, et al: Phosphorylation of amyloid precursor protein by mutant LRRK2 promotes AICD activity and neurotoxicity in Parkinson’s disease. Sci Signal 2017, 10:eaam6790. 10.1126/scisignal.aam6790

21. Zhang ZW, Tu H, Jiang M, Vanan S, Chia SY, Jang SE, Saw WT, Ong ZW, Ma DR, Zhou ZD, et al: The APP intracellular domain promotes LRRK2 expression to enable feed-forward neurodegenerative mechanisms in Parkinson’s disease. Sci Signal 2022, 15:eabk3411. 10.1126/scisignal.abk3411

22. Branca C, Sarnico I, Ruotolo R, Lanzillotta A, Viscomi AR, Benarese M, Porrini V, Lorenzini L, Calza L, Imbimbo BP, et al: Pharmacological targeting of the beta-amyloid precursor protein intracellular domain. Sci Rep 2014, 4:4618. 10.1038/srep04618

23. Hughes JP, Rees S, Kalindjian SB, Philpott KL: Principles of early drug discovery. Br J Pharmacol 2011, 162:1239–1249. 10.1111/j.1476-5381.2010.01127.x

24. Nair AB, Jacob S: A simple practice guide for dose conversion between animals and human. J Basic Clin Pharm 2016, 7:27–31. 10.4103/0976-0105.177703

25. Cummings JL, Zhou Y, Van Stone A, Cammann D, Tonegawa-Kuji R, Fonseca J, Cheng F: Drug repurposing for Alzheimer’s disease and other neurodegenerative disorders. Nat Commun 2025, 16:1755. 10.1038/s41467-025-56690-4

26. Hatami S, Sirous H, Mahnam K, Najafipour A, Fassihi A: Preparing a database of corrected protein structures important in cell signaling pathways. Res Pharm Sci 2023, 18:67–77. 10.4103/1735-5362.363597

27. Roos K, Wu C, Damm W, Reboul M, Stevenson JM, Lu C, Dahlgren MK, Mondal S, Chen W, Wang L, et al: OPLS3e: Extending Force Field Coverage for Drug-Like Small Molecules. J Chem Theory Comput 2019, 15:1863–1874. 10.1021/acs.jctc.8b01026

28. Friesner RA, Banks JL, Murphy RB, Halgren TA, Klicic JJ, Mainz DT, Repasky MP, Knoll EH, Shelley M, Perry JK, et al: Glide:D A New Approach for Rapid, Accurate Docking and Scoring. 1. Method and Assessment of Docking Accuracy. Journal of Medicinal Chemistry 2004, 47:1739–1749. 10.1021/jm0306430

29. Genheden S, Ryde U: The MM/PBSA and MM/GBSA methods to estimate ligand-binding affinities. Expert Opin Drug Discov 2015, 10:449–461. 10.1517/17460441.2015.1032936

30. Tu H, Yeo XY, Zhang ZW, Zhou W, Tan JY, Chi L, Chia SY, Li Z, Sim AY, Singh BK, et al: NOTCH2NLC GGC intermediate repeat with serine induces hypermyelination and early Parkinson’s disease-like phenotypes in mice. Mol Neurodegener 2024, 19:91. 10.1186/s13024-024-00780-2

31. Tu H, Zhang ZW, Qiu L, Lin Y, Jiang M, Chia SY, Wei Y, Ng ASL, Reynolds R, Tan EK, Zeng L: Increased expression of pathological markers in Parkinson’s disease dementia post-mortem brains compared to dementia with Lewy bodies. BMC Neurosci 2022, 23:3. 10.1186/s12868-021-00687-4

32. Ashburn TT, Thor KB: Drug repositioning: identifying and developing new uses for existing drugs. Nat Rev Drug Discov 2004, 3:673–683. 10.1038/nrd1468

33. Cummings J, Lee G, Ritter A, Sabbagh M, Zhong K: Alzheimer’s disease drug development pipeline: 2020. Alzheimers Dement (N Y) 2020, 6:e12050. 10.1002/trc2.12050

34. Li X, Wang QJ, Pan N, Lee S, Zhao Y, Chait BT, Yue Z: Phosphorylation-dependent 14-3-3 binding to LRRK2 is impaired by common mutations of familial Parkinson’s disease. PLoS One 2011, 6:e17153. 10.1371/journal.pone.0017153

35. Muda K, Bertinetti D, Gesellchen F, Hermann JS, von Zweydorf F, Geerlof A, Jacob A, Ueffing M, Gloeckner CJ, Herberg FW: Parkinson-related LRRK2 mutation R1441C/G/H impairs PKA phosphorylation of LRRK2 and disrupts its interaction with 14-3-3. Proc Natl Acad Sci U S A 2014, 111:E34–43. 10.1073/pnas.1312701111

36. Cookson MR: The role of leucine-rich repeat kinase 2 (LRRK2) in Parkinson’s disease. Nat Rev Neurosci 2010, 11:791–797. 10.1038/nrn2935

37. Zhu H, Hixson P, Ma W, Sun J: Pharmacology of LRRK2 with type I and II kinase inhibitors revealed by cryo-EM. Cell Discov 2024, 10:10. 10.1038/s41421-023-00639-8

38. Kumar BH, Manandhar S, Mehta CH, Nayak UY, Pai KSR: Structure-based docking, pharmacokinetic evaluation, and molecular dynamics-guided evaluation of traditional formulation against SARS-CoV-2 spike protein receptor bind domain and ACE2 receptor complex. Chem Zvesti 2022, 76:1063–1083. 10.1007/s11696-021-01917-z

39. Li Q, Li S, Fang J, Yang C, Zhao X, Wang Q, Zhou W, Zheng W: Artemisinin Confers Neuroprotection against 6-OHDA-Induced Neuronal Injury In Vitro and In Vivo through Activation of the ERK1/2 Pathway. Molecules 2023, 28:5527.

40. Marriott JJ, Miyasaki JM, Gronseth G, O’Connor PW, Therapeutics, Technology Assessment Subcommittee of the American Academy of N: Evidence Report: The efficacy and safety of mitoxantrone (Novantrone) in the treatment of multiple sclerosis [RETIRED]: Report of the Therapeutics and Technology Assessment Subcommittee of the American Academy of Neurology. Neurology 2010, 74:1463–1470. 10.1212/WNL.0b013e3181dc1ae0

41. Gonsette RE: Mitoxantrone in progressive multiple sclerosis: when and how to treat? J Neurol Sci 2003, 206:203–208. 10.1016/s0022-510x(02)00335-0

42. Osborne CK, Drelichman A, Von Hoff DD, Crawford ED: Mitoxantrone: modest activity in a phase II trial in advanced prostate cancer. Cancer Treat Rep 1983, 67:1133–1135.

43. Nair A, Morsy MA, Jacob S: Dose translation between laboratory animals and human in preclinical and clinical phases of drug development. Drug Dev Res 2018, 79:373–382. 10.1002/ddr.21461

44. Yue M, Hinkle KM, Davies P, Trushina E, Fiesel FC, Christenson TA, Schroeder AS, Zhang L, Bowles E, Behrouz B, et al: Progressive dopaminergic alterations and mitochondrial abnormalities in LRRK2 G2019S knock-in mice. Neurobiol Dis 2015, 78:172–195. 10.1016/j.nbd.2015.02.031

45. Xenias HS, Chen C, Kang S, Cherian S, Situ X, Shanmugasundaram B, Liu G, Scesa G, Chan CS, Parisiadou L: R1441C and G2019S LRRK2 knockin mice have distinct striatal molecular, physiological, and behavioral alterations. Commun Biol 2022, 5:1211. 10.1038/s42003-022-04136-8

46. Chang EES, Ho PW, Liu HF, Pang SY, Leung CT, Malki Y, Choi ZY, Ramsden DB, Ho SL: LRRK2 mutant knock-in mouse models: therapeutic relevance in Parkinson’s disease. Transl Neurodegener 2022, 11:10. 10.1186/s40035-022-00285-2

47. Avasarala JR, Cross AH, Clifford DB, Singer BA, Siegel BA, Abbey EE: Rapid onset mitoxantrone-induced cardiotoxicity in secondary progressive multiple sclerosis. Mult Scler 2003, 9:59–62. 10.1191/1352458503ms896oa

48. Fleischer V, Salmen A, Kollar S, Weyer V, Siffrin V, Chan A, Zipp F, Luessi F: Cardiotoxicity of mitoxantrone treatment in a german cohort of 639 multiple sclerosis patients. J Clin Neurol 2014, 10:289–295. 10.3988/jcn.2014.10.4.289

49. Shaikh AY, Suryadevara S, Tripathi A, Ahmed M, Kane JL, Escobar J, Cerny J, Nath R, McManus DD, Shih J, et al: Mitoxantrone-Induced Cardiotoxicity in Acute Myeloid Leukemia-A Velocity Vector Imaging Analysis. Echocardiography 2016, 33:1166–1177. 10.1111/echo.13245

50. Alqahtani MS, Kazi M, Alsenaidy MA, Ahmad MZ: Advances in Oral Drug Delivery. Front Pharmacol 2021, 12:618411. 10.3389/fphar.2021.618411

51. Homayun B, Lin X, Choi HJ: Challenges and Recent Progress in Oral Drug Delivery Systems for Biopharmaceuticals. Pharmaceutics 2019, 11:129. 10.3390/pharmaceutics11030129

52. Cohen BA, Mikol DD: Mitoxantrone treatment of multiple sclerosis: safety considerations. Neurology 2004, 63:S28–32. 10.1212/wnl.63.12_suppl_6.s28

53. Crossley RJ: Clinical safety and tolerance of mitoxantrone. Semin Oncol 1984, 11:54–58.

54. Reis-Mendes A, Dores-Sousa JL, Padrao AI, Duarte-Araujo M, Duarte JA, Seabra V, Goncalves-Monteiro S, Remiao F, Carvalho F, Sousa E, et al: Inflammation as a Possible Trigger for Mitoxantrone-Induced Cardiotoxicity: An In Vivo Study in Adult and Infant Mice. Pharmaceuticals (Basel) 2021, 14. 10.3390/ph14060510

55. Dores-Sousa JL, Duarte JA, Seabra V, Bastos Mde L, Carvalho F, Costa VM: The age factor for mitoxantrone’s cardiotoxicity: multiple doses render the adult mouse heart more susceptible to injury. Toxicology 2015, 329:106–119. 10.1016/j.tox.2015.01.006

56. Seiter K: Toxicity of the topoisomerase II inhibitors. Expert Opin Drug Saf 2005, 4:219–234. 10.1517/14740338.4.2.219

57. Lazic SE, Semenova E, Williams DP: Determining organ weight toxicity with Bayesian causal models: Improving on the analysis of relative organ weights. Sci Rep 2020, 10:6625. 10.1038/s41598-020-63465-y

58. Sellers RS, Morton D, Michael B, Roome N, Johnson JK, Yano BL, Perry R, Schafer K: Society of Toxicologic Pathology position paper: organ weight recommendations for toxicology studies. Toxicol Pathol 2007, 35:751–755. 10.1080/01926230701595300

59. Rossato LG, Costa VM, Dallegrave E, Arbo M, Silva R, Ferreira R, Amado F, Dinis-Oliveira RJ, Duarte JA, de Lourdes Bastos M, et al: Mitochondrial cumulative damage induced by mitoxantrone: late onset cardiac energetic impairment. Cardiovasc Toxicol 2014, 14:30–40. 10.1007/s12012-013-9230-2

60. Bieber JM, Sanman LE, Sun X, Hammerlindl H, Bao F, Roth MA, Koleske ML, Huang L, Aweeka F, Wu LF, Altschuler SJ: Differential toxicity to murine small and large intestinal epithelium induced by oncology drugs. Commun Biol 2022, 5:99. 10.1038/s42003-022-03048-x

61. Wang J, Cai E, An X, Wang J: Ginaton reduces M1-polarized macrophages in hypertensive cardiac remodeling via NF-kappaB signaling. Front Pharmacol 2023, 14:1104871. 10.3389/fphar.2023.1104871

62. Liu W, Sun J, Guo Y, Liu N, Ding X, Zhang X, Chi J, Kang N, Liu Y, Yin X: Calhex231 ameliorates myocardial fibrosis post myocardial infarction in rats through the autophagy-NLRP3 inflammasome pathway in macrophages. J Cell Mol Med 2020, 24:13440–13453. 10.1111/jcmm.15969

63. Hartung HP, Gonsette R, Konig N, Kwiecinski H, Guseo A, Morrissey SP, Krapf H, Zwingers T, Mitoxantrone in Multiple Sclerosis Study G: Mitoxantrone in progressive multiple sclerosis: a placebo-controlled, double-blind, randomised, multicentre trial. Lancet 2002, 360:2018–2025. 10.1016/S0140-6736(02)12023-X

64. Scott LJ, Figgitt DP: Mitoxantrone: a review of its use in multiple sclerosis. CNS Drugs 2004, 18:379–396. 10.2165/00023210-200418060-00010

65. Aarsland D, Zaccai J, Brayne C: A systematic review of prevalence studies of dementia in Parkinson’s disease. Mov Disord 2005, 20:1255–1263. 10.1002/mds.20527

66. Aarsland D, Bernadotte A: Epidemiology of dementia associated with Parkinson’s disease. In Cognitive Impairment and Dementia in Parkinson’s Disease. Edited by Emre M: Oxford University Press; 2015: 5–16 10.1093/med/9780199681648.003.0002

67. Kane JPM, Surendranathan A, Bentley A, Barker SAH, Taylor JP, Thomas AJ, Allan LM, McNally RJ, James PW, McKeith IG, et al: Clinical prevalence of Lewy body dementia. Alzheimers Res Ther 2018, 10:19. 10.1186/s13195-018-0350-6

68. Kotzbauer PT, Trojanowsk JQ, Lee VM: Lewy body pathology in Alzheimer’s disease. J Mol Neurosci 2001, 17:225–232. 10.1385/jmn:17:2:225

69. Eleuteri S, Di Giovanni S, Rockenstein E, Mante M, Adame A, Trejo M, Wrasidlo W, Wu F, Fraering PC, Masliah E, Lashuel HA: Novel therapeutic strategy for neurodegeneration by blocking Abeta seeding mediated aggregation in models of Alzheimer’s disease. Neurobiol Dis 2015, 74:144–157. 10.1016/j.nbd.2014.08.017

70. Kingwell E, Koch M, Leung B, Isserow S, Geddes J, Rieckmann P, Tremlett H: Cardiotoxicity and other adverse events associated with mitoxantrone treatment for MS. Neurology 2010, 74:1822–1826. 10.1212/WNL.0b013e3181e0f7e6

71. Hamzehloo A, Etemadifar M: Mitoxantrone-induced cardiotoxicity in patients with multiple sclerosis. Arch Iran Med 2006, 9:111–114.

72. Potashkin JA, Blume SR, Runkle NK: Limitations of animal models of Parkinson’s disease. Parkinsons Dis 2010, 2011:658083. 10.4061/2011/658083

73. Antony PM, Diederich NJ, Balling R: Parkinson’s disease mouse models in translational research. Mamm Genome 2011, 22:401–419. 10.1007/s00335-011-9330-x

74. Gilsbach BK, Messias AC, Ito G, Sattler M, Alessi DR, Wittinghofer A, Kortholt A: Structural Characterization of LRRK2 Inhibitors. J Med Chem 2015, 58:3751–3756. 10.1021/jm5018779

